# *Ixodes ricinus* bites promote allergic skin inflammation and intestinal tuft and mast cell expansion in mice

**DOI:** 10.1101/2024.07.02.600632

**Authors:** Juan Manuel Leyva-Castillo, Maria Strakosha, Sophia E.M. Smith, Daniela Vega-Mendoza, Megan Elkins, Janet Chou, Peter Vogel, Nathalie Boulanger

## Abstract

**Background:** Tick bites often promote local allergic reactions in the skin and predispose to red meat allergy. The mechanisms involved in these processes are not fully understood. Here we investigated the local changes to the skin and intestine induced by tick bites.

**Methods:** C3H/HEN or Balb/c mice were subjected to either tick bites by *Ixodes ricinus* (*I. ricinus*) or mechanical skin injury. Skin or intestine was analyzed a different time point by transcriptomic and histological techniques.

**Results:** Our results indicate that *I. ricinus* bites promote epidermal hyperplasia, spongiosis and an accumulation of eosinophils and mast cells in the bitten skin. In addition, *I. ricinus* bites promote the expression of genes and activate pathways also induced by mechanical skin injury elicited by tape stripping. Remarkably, similar to tape stripping, *I. ricinus* bites promote an increase in total serum IgE, and intestinal tuft cell and mast cell expansion.

**Conclusion:** *I. ricinus* bites in mice promote cutaneous inflammation that resembles allergic skin inflammation, as well as intestinal changes that could play a role in the predisposition to red meat allergy.

## INTRODUCTION

Ticks are vectors for numerous infectious diseases, such as Lyme disease, anaplasmosis, rocky mountain spotted fever and tularemia (1). Tick bites cause mild allergic reactions, which manifest as a cutaneous rash, promoting the accumulation of eosinophils, basophils, T cells and the degranulation of mast cells in the affected area (1-4). In addition, tick bites are one of the major predisposing factors to red meat allergy (5-8). Red meat allergy is a unique food allergy characterized by delayed hypersensitivity to red meat mediated by an IgE antibody response against alpha-galactose (gal) moieties (5-8). Two different phases could be described in this disease, the first one, the sensitization phase, is characterized by the presence of IgE-specific against alpha-gal moieties, commonly associated with tick bites and the second one, the effector phase, which is elicited following the consumption of red meat (5-8). How tick bites promote skin inflammation and predispose to red meat allergy is not completely understood.

Tick bites or tick salivary products promote a type 2 dominated immune response in mice, including an increased number of Th2 cells in the skin draining lymph node and spleen, increased levels of IL-4 in skin draining lymph nodes, and increased levels of serum total IgE (9-15). Moreover, tick salivary products inhibited Th1 polarization and the production of IFNγ by skin draining lymph node cells (9, 11). Similar to tick bites, mechanical skin injury induced by tape stripping polarizes the immune response to a type 2 dominated phenotype, including increased production of Th2 cytokines and increased levels of total IgE (16-18). Tape stripping and topical application of antigen elicits allergic skin inflammation, characterized by increase epidermal thickness, accumulation of eosinophils and mast cells, and upregulation of cutaneous *Il4* and *Il13* (17, 19, 20). In addition, mechanical skin injury promotes increase of total serum IgE and the expansion of intestinal mast cells, which promote more severe passive and active anaphylaxis (21). Whether mechanical skin injury and tick bites triggered similar mechanisms have not been evaluated.

## Materials and Methods

### Mice

C3H/HEN and Balb/c mice were obtained from Charles River. All mice were kept in a pathogen-free environment. All procedures were performed in accordance with the Animal Care and Use Committee of the Institut de bactériologie, Universite de Strasbourg and Boston Children’s Hospital.

### *I. ricinus* infestation

Ticks were collected on vegetation in forested areas in Alsace (France) and tested them for the presence of *Borrelia burgdorferi* sensu lato or *Anaplasma phagocytophilum* as described (22). We also used *I. ricinus* from a laboratory reared-tick colony but the effect on the jejunum was absent (data not shown). After shaving the back of C3H/HeN mouse, a plastic cap was fixed with wax to the mouse’s back. Five female ticks or 10 nymphs per mouse were introduced into the cap which was sealed. For the control animals, plastic cap was just fixed to the back of the animal. After 4 days of blood meal (complete blood meal for the nymphs), the cap was removed, and all ticks were collected. At day 4, 10 and 14 days, the skin at the site of the tick bite and the jejunum were removed. For the control animals, the cap was removed, and skin biopsy and jejunum were collected as for the animals bitten by ticks.

### Mechanical skin injury induced by tape stripping

6- to 8-week-old female Balb/c mice were anesthetized; their back skin was shaved and tape-stripped with a film dressing (Tegaderm TM, 3M) 6 times. Skin samples were collected 6 h after tape stripping and keep in RNA later.

### Skin histology

Skin and gut specimens were fixed in 4% paraformaldehyde embedded in paraffin. Skin sections (5μm) were stained with hematoxylin/eosin (H&E) and toluidine blue. ImageJ was used for the quantification of the epidermal thickness and mast cells as previously described(17).

### Immunohistochemical (IHC) staining

IHC staining of MBP (for eosinophils) was done as previously described (19). Paraffin sections were treated with 0.6% H_2_O_2_ to block the endogenous peroxidase activity, followed by digestion with Pepsin solution (Invitrogen) to retrieve antigen. Slides were then blocked with rabbit serum (Vector Laboratories) and incubated with rat-anti-mouse MBP (provided by Dr. Elizabeth Jacobsen, Mayo Clinic, Rochester, USA). After incubation with biotinylated rabbit anti-rat IgG, followed by AB complex (Vector Laboratories), staining was finally visualized with AEC+ high sensitivity substrate chromogen solution (Vector Laboratories) and counterstained with hematoxylin. IHC staining of MCPT1, MCPT4 and DCAMKL1 was done as previously described (23). Gut sections (5μm) underwent antigen retrieval at 100°C for 20 min in Epitope Retrieval solution 2 (ER2) on a Bond Max immunostainer (Leica Biosystems, Bannockburn, IL). A rat monoclonal primary antibody to mast cell protease 1 (MCPT1;Cat# 14-5503-82; eBiosciences, San Diego, CA) diluted 1:30 was applied for 15 minutes followed by biotinylated secondary rabbit anti-rat antibody diluted 1:400 (BA-4001; Vector Laboratories, Burlingame, CA) for 10 minutes and then ready-to-use Bond polymer refine detection kit with 3,3’- diaminobenzidine (DAB) as chromogenic substrate and a hematoxylin counterstain (DS9800; Leica Biosystems). For detection of lpMMCs and tuft cells, tissue sections underwent antigen retrieval in a prediluted Cell Conditioning Solution (*CC1*) (Ventana Medical Systems, Indianapolis, IN) for 32 min. For lpMMCs, a goat polyclonal antibody to mast cell protease 4 (MCPT4; LS-B5958; LifeSpan Biosciences, Seattle, WA) was diluted 1:500 and applied for 30 min at RT followed by biotinylated secondary rabbit anti-goat antibody 1:400 (BA-5000; Vector Laboratories) for 10 minutes. For detection of tuft cells, a rabbit polyclonal antibody to doublecortin like kinase 1 (DCAMKL1; ab31704; Abcam, Cambridge, MA) diluted 1:1,000 was applied for 30 min at RT followed directly by the OmniMap anti-Rabbit HRP kit (Ventana Medical Systems) for detection, with Discovery Purple as chromogen for lpMMCs and ChromoMap DAB for tuft cells (both from Ventana Medical Systems).

### Analysis of cytokine expression in skin and jejuna

Total skin RNA was extracted with RNeasy Mini Kit (Qiagen). cDNA was prepared with iScript cDNA synthesis kit (Biorad). PCR reactions were run on Quantstudio 5 (Applied Biosystems) sequence detection system platform. Taqman primers and probes were obtained from Life technologies. The housekeeping gene β_2_-microglobulin was used as an internal control. Relative mRNA expression was quantified using the 2^-ΔΔCt^ method.

### Transcriptomic analysis

Total skin RNA was isolated as described above, followed by cDNA synthesis using the SuperScript VILO cDNA Synthesis Kit (ThermoFisher Scientific). The Ion AmpliSeq Transcriptome Mouse Gene Expression Kit was used to prepare bar-coded libraries and sequenced by using an Ion S5 next-generation sequencer. The AmpliSeqRNA plug-in (ThermoFisher Scientific) was used to calculate differential gene expression analysis. Pathway analysis was performed by using Ingenuity Pathway Analysis (Qiagen) on genes with at least a 2.0-fold difference between the conditions (p<.05).

### Serum IgE Determination

Preimmune sera were first collected and then after tick bites, immune sera were collected at day 14. Microtiter plates (R&D Systems) were first coated with a monoclonal rat antimouse IgE (clone R35–72; BD Biosciences), then, coated plates were blocked with PBS–3% rat serum and incubated with serum samples and with a biotinylated rat antimouse IgE (clone R35–118; BD Biosciences). Avidin-HRP (Ebiosciences) and TMB (tetramethylbenzidine) substrate reagent (Ebiosciences) were used for detection. Serum total IgE concentration was determined from a standard curve of mouse IgE (clone C38–2; BD Biosciences).

### Statistical analysis

Statistical significance was determined by the two-tailed Student’s t test. A p value <0.05 was considered statistically significant.

## RESULTS

### Tick bites from *Ixodes ricinus* promote skin inflammation resembling atopic dermatitis

To investigate the changes in the skin induced by tick bites, we analyzed both the intact skin and *Ixodes ricinus* (*I. ricinus*) bitten skin of C3H/HEN mice by histology. Following blood feeding for 4 days, all ticks were removed from the back skin and skin was evaluated immediately after and following 6 (day 10) or 10 (day 14) days as depicted in **figure 1A**. We analyzed the skin of bitten mice for the presence of *B. burgdorferi* sensu lato and *A. phagocytophilum* by PCR in the experiments carried with field-collected ticks. We detected *Borrelia* and *Anaplasma* in several but not all the bitten skin (Supplementary table 1). After removing *I. ricinus*, the back skin looked comparable to unbitten skin, however at day 10 the back skin of bitten mice showed increased epidermal and dermal thickness, spongiosis and the accumulation of mononuclear cells, which is decreased almost to the levels seen in intact skin at day 14 (**Fig. 1B**). Bitten skin at day 10 exhibited increased accumulation of mast cells in the dermis and eosinophil (major basic protein [MBP]^+^ cells) accumulation into the hypodermis (**Fig. 1C and D**). These results suggest that tick bites promote skin inflammation that histologically resemble allergic skin inflammation.

**Figure 1.**
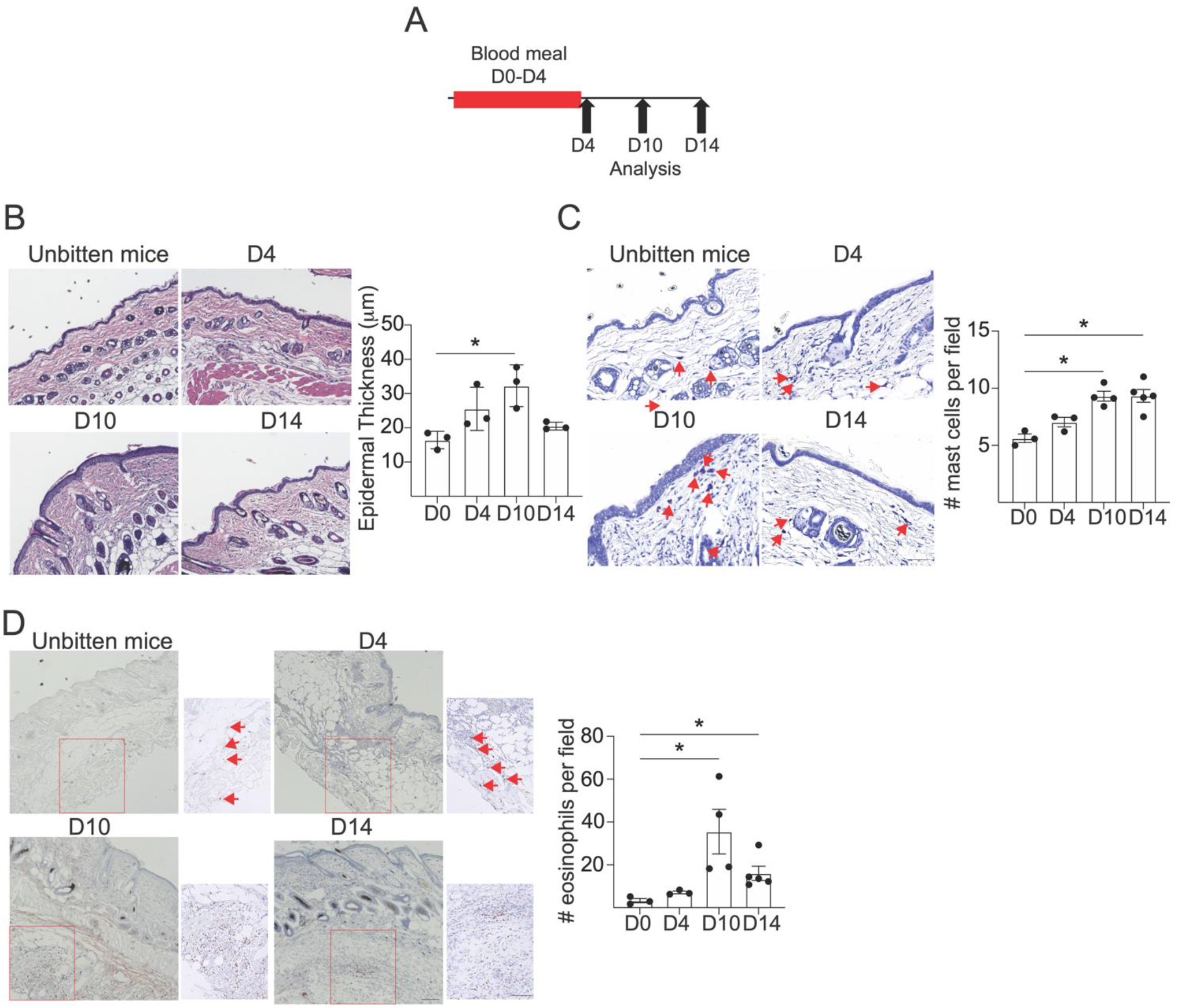
Tick bites from *Ixodes ricinus* promote skin inflammation resembling atopic dermatitis. **A**. Experimental protocol. **B-D**. Representative H&E staining and epidermal thickness quantitation (B), representative toluidine blue staining and mast cell quantitation (C) and representative Major basic protein (MBP) immunohistochemistry staining and eosinophils quantitation (D) in unbitten skin from C3H/HEN mice or subjected to *I. ricinus* bites at different time points. Red arrows indicate toluidine blue+ (Mast cells) and MBP+ cells (eosinophils) *p<0.05 by t test (one experiment with a n=3-5 mice per experiment).

### The gene expression profile of skin subjected to tick bites from *I. ricinus* resembles skin subjected to mechanical skin injury induced by tape stripping

To gain insight in the potential mechanisms induced by tick bites in the skin of the host we performed transcriptomic analysis on the skin at day 4. Bitten skin exhibited 588 genes that were differentially expressed by more than 2-fold (p<0.05) compared to unbitten skin. Of these 588 genes, 207 were downregulated and 381 were upregulated (**Fig. 2A** and supplementary table 2). Ingenuity pathway analysis (IPA) canonical pathway analysis of genes differentially expressed between bitten skin and unbitten skin revealed that tick bites induce the expression of genes with roles in granulocyte and agranulocyte adhesion and diapedesis, pathogen recognition, neutrophil and macrophage activation, and IL-17, IL-33, and IL-13 signaling among others (**Fig. 2B**). In addition, IPA upstream analysis identified inflammatory cytokines such as TNFα, IL-1α, IL-1β, the Th17 cytokine IL-17A and the Th2 cytokine IL-33, as well as the toll like receptors TLR3, TLR4 and TLR9 and the adaptor molecule MYD88 (**Fig. 2C**). We have previously shown that mechanical skin injury induced by tape stripping promotes the production of type 2 and type 17 cytokines (21, 24-26). To investigate whether tick bites from *I. ricinus* promote a similar gene expression profile as mechanical skin injury induced by tape stripping, we performed transcriptomic analysis on shave and tape stripped skin from WT Balb/c mice. Tape stripped skin exhibited 2576 genes that were differentially expressed by more than 2-fold (p<0.05) compared to shaved skin. Of these 2576 genes, 1615 were downregulated and 961 were upregulated (**Fig. 2D** and supplementary table 3). Similar to skin subjected to *I. ricinus* bites, IPA canonical pathway analysis of genes differentially expressed between shaved skin and tape stripped skin revealed that mechanical skin injury induces the expression of genes with roles in granulocyte and agranulocyte adhesion and diapedesis, pathogen recognition, neutrophil and macrophage activation, and IL-17, IL-33 and IL-13 signaling (**Fig 2E**). In addition, IPA upstream analysis identified inflammatory cytokines such as TNFα, IL-1α, IL-1β, the Th17 cytokine IL-17A and the Th2 cytokine IL-33, the toll like receptors TLR3, TLR4 and TLR9, and the adaptor molecule MYD88 (**Fig. 2F**). To investigate whether tick bites and mechanical skin injury induced by tape stripping share gene expression profiles, we compared genes differentially expressed by more than 2-fold (p<0.05) in bitten skin at day 4 and unbitten skin; along with those of tape stripped skin and shaved skin. We identified 315 genes that were shared in the two datasets, of these 315 genes,184 were upregulated in both datasets, 113 genes were downregulated in both datasets and 18 were differentially regulated (**Fig. 2G**). The expression of *Cd180, Ncr1, Arg1, Clec4a, C5ar1* and *Cd163*, related with defense response, were increased in the skin of bitten mice but decreased in tape stripped skin. Remarkably, comparative analysis of pathways canonical pathway analysis between of datasets showed that 110 pathways were shared by our datasets. Using only the pathways with a predicted activation or inhibition by IPA (Z-score ± 2), our analysis showed that 22 pathways were activated or inhibited similarly between our datasets (**Fig. 2H**). Our results indicate that tick bites promote a gene expression profile in the skin that resembles the one induced by mechanical skin injury.

**Figure 2.**
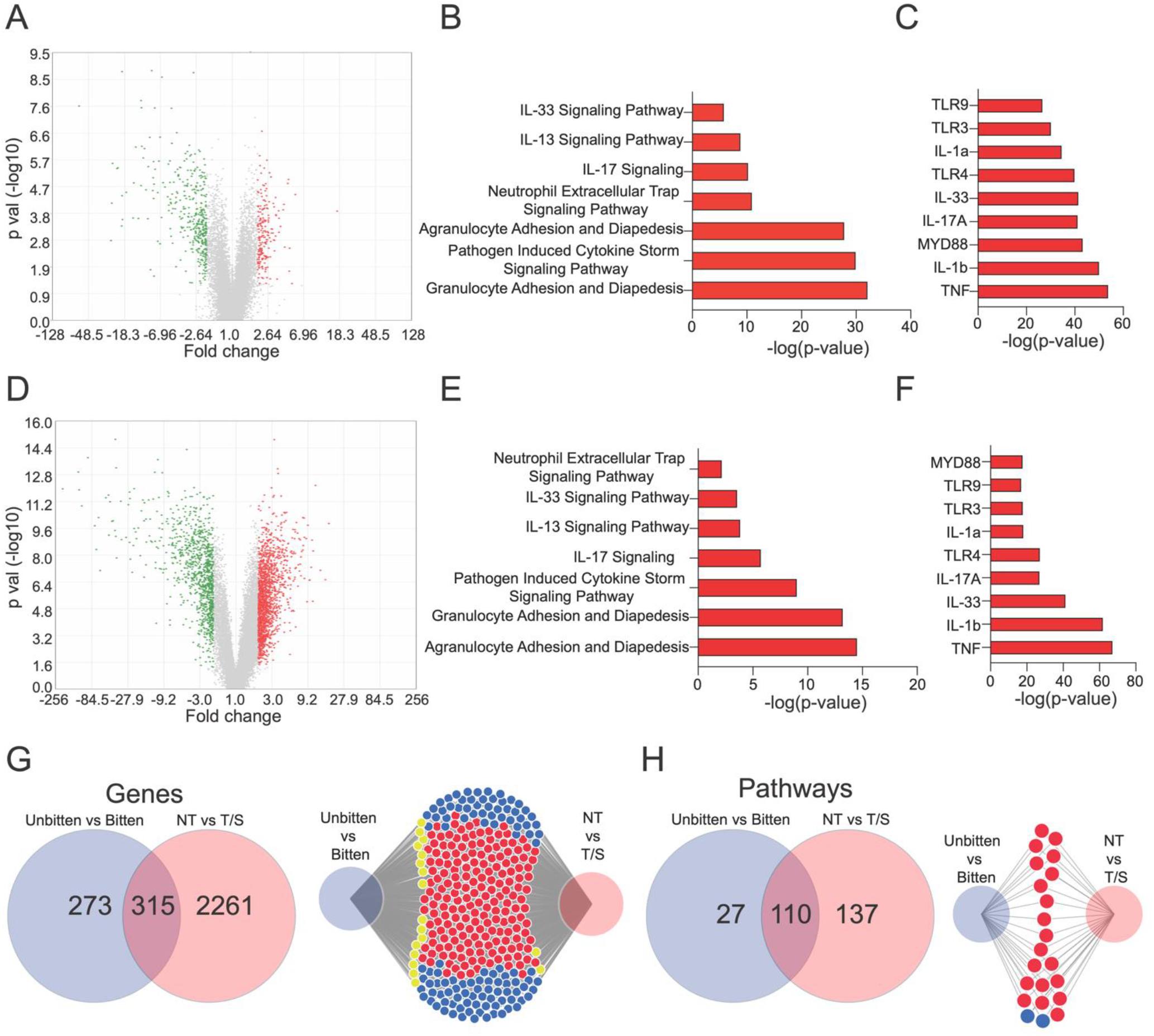
Gene expression profile from skin subjected to tick bites from *Ixodes ricinus* resembles skin subjected to mechanical skin injury induced by tape stripping. **A-C**. Volcano plot (A) bar plots for enriched Ingenuity pathway analysis (IPA) canonical pathway analysis (A) or upstream regulator analysis (C) of differentially expressed genes by at least 2.0-fold difference (p<0.05). of unbitten skin from C3H/HEN mice and skin subjected to *I. ricinus* bites at day 4. **D-F**. Volcano plot (A) bar plots for enriched IPA canonical pathway analysis (A) or upstream regulator analysis (C) of differentially expressed genes by at least 2.0-fold difference (p<0.05). of shaved skin from Balb/c mice and skin subjected to tape stripping at 6 h. **G-H**. Venn and Divenn diagrams displaying differentially expressed genes (G) or pathways (H) in *I*. *ricinus* bitten skin and unbitten skin from C3H/HEN mice and tape stripped and shaved skin from Balb/c mice datasets using DiVenn. Blue and red circles denote genes down- regulated and upregulated, respectively. Yellow nodes denote differences between datasets. Data presented in this figure is coming from 2 independent experiments.

### Tick bites from *I. ricinus* promote a type 2-regulated gene expression profile in the skin and an increase in total IgE levels

To investigate whether tick bites promote a type 2 immune response, similar to allergic skin inflammation at day 10, we evaluated *Il4* and *Il13* mRNA levels at day 10. Remarkably, the levels of *Il4* and *Il13* mRNA were increased in bitten skin compared with unbitten skin (**Fig. 3A**). In addition, bitten skin showed increased *Ifng* mRNA levels, however *Il17a* was not detected in bitten or unbitten skin at day 10 (**Fig. 3A** and data not shown). Consistent with these results, IPA upstream analysis of differentially expressed genes between unbitten skin and bitten skin at day 10 identified several genes controlled by IL-4 and/or IL-13 (**Fig. 3B-D** and supplementary table 4). Our analysis inferred that IL-4 and IL-13 upregulate the expression of genes coding for matrix metalloproteinases (*Mmp10, Mmp12* and *Mmp13*), collagen 1 (*Col1a1* and *Col1a2*), related with NK cells (*Gzma, Gzmb* and *Il7r*) and related with alternative activated macrophage (Arg2, *Retnlg and fn1*). Moreover IL-4 and IL-13 suppressed the expression of *Il12rb*, and genes related with inflammatory macrophages *(Fabp4, Alox5, Gpx3, Pparg* and *Ptges*) among others (**Fig. 3B-D**).

**Figure 3.**
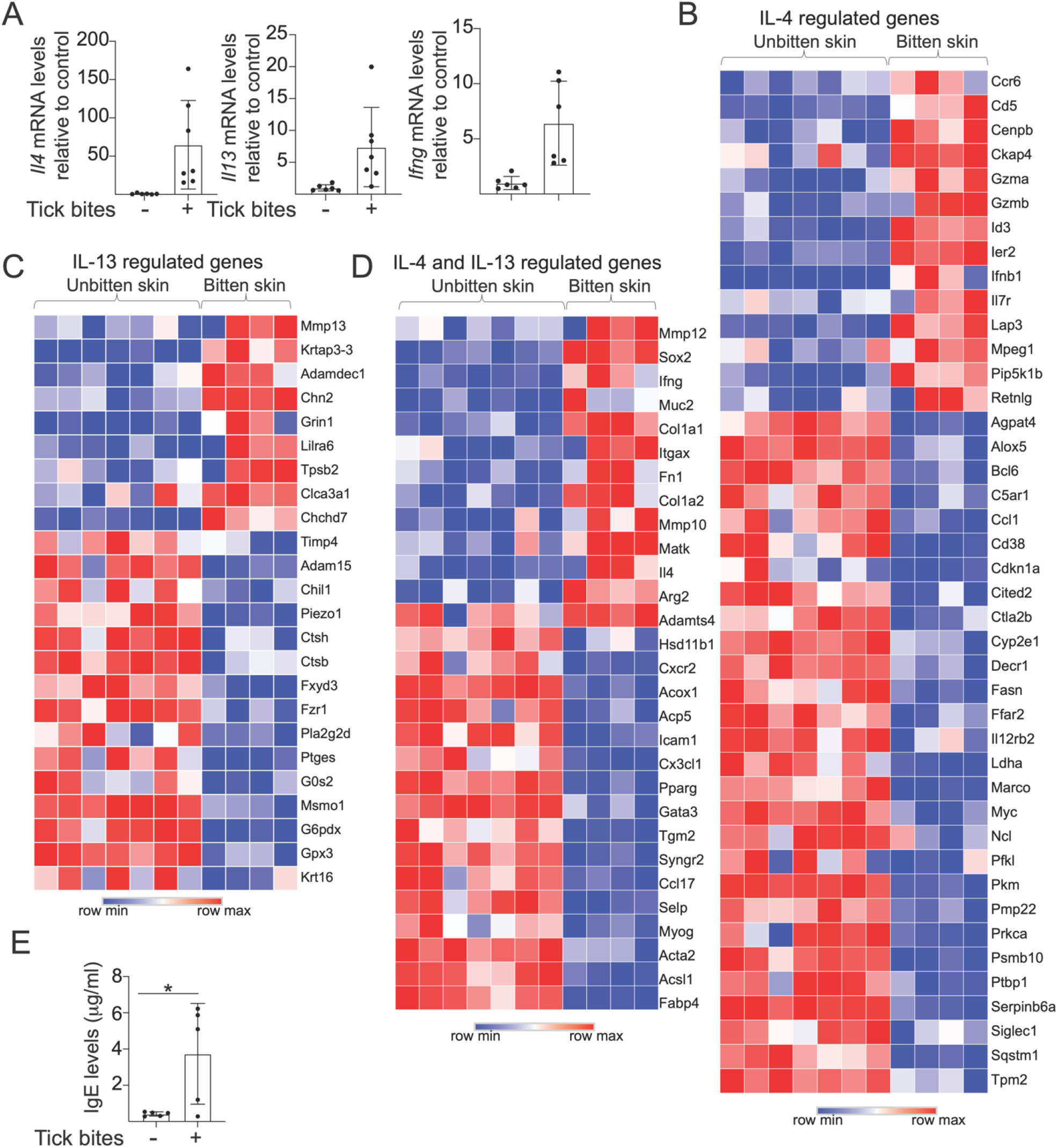
Tick bites from *Ixodes ricinus* promote a type 2-regulated gene expression profile in the skin and increase in total IgE serum levels. **A**. Il4 (left), Il13 (middle) and Ifng (rigth) mRNA levels in unbitten skin from C3H/HEN mice or subjected to I. ricinus bites at day 10. **B-D**. Heat map of inferred genes regulated by IL-4 (B), IL-13 (C) or both (D) by Ingenuity pathway analysis upstream regulator analysis of differentially expressed genes by at least 2.0-fold difference (p<0.05). **E**. Total serum IgE levels in C3H/HEN mice or subjected to *I. ricinus* bites at day 14. Data presented in panel A-D is coming from 2 independent experiments, n=3-5 mice per experiment. Data presented in panel E is coming from 1 experiment with 5 mice per group.* p<0.05 by paired t test.

Allergic skin inflammation is often accompanied by the presence of allergen-specific IgE and increased total IgE levels in circulation (17, 19, 21). To investigate whether tick bites promote the production of IgE, we measured IgE serum levels in C3H/HEN mice before and following *I. ricinus* infestation. Our results showed that following *I. ricinus* bites C3H/HEN mice increased total IgE serum levels (**Fig. 3E**). We were unable to measure alpha-gal-IgE as mice, in contrast to human and apes, express a fully functional α-1,3-galactosyltransferase and murine proteins abundantly carry α-Gal residues (27). Our results suggest that tick bites promote inflammation that resembles atopic dermatitis.

### Tick bites from *I. ricinus* promote increases in the number of tuft cells and mast cells in the jejunum

To investigate whether tick bites from *I. ricinus* provoke changes in the jejunum, we analyzed the jejunum of bitten mice at day 10. Jejuna from bitten mice exhibited increased *Dclk1* and *Il25* mRNA expression compared with unmanipulated mice (**Fig. 4A**). Consistent with this, immunohistochemistry analysis confirmed an increased number of DCLK1^+^ cells in the jejuna of bitten mice (**Fig. 4B**). In addition, transcriptomic analysis of jejunum at day 10 showed an increase in the expression of mast cell-related genes, including *Mcpt1, Mcpt2, Fcer1a, Ms4a2, Hpgd* and *Ptgr1* in bitten mice compared with unbitten mice (**Fig. 4C** and supplementary table 5). In line with this, immunohistochemistry analysis confirmed an increased numbers of mucosal mast cells (MCPT1^+^) and the appearance of connective tissue mast cells (MCPT4^+^) in the jejuna of bitten mice compared with unbitten mice (**Fig. 4D-E**). Our results indicate that tick bites from *I. ricinus* promote an increase in tuft cells and mast cells in the jejunum of C3H/HEN mice.

**Figure 4.**
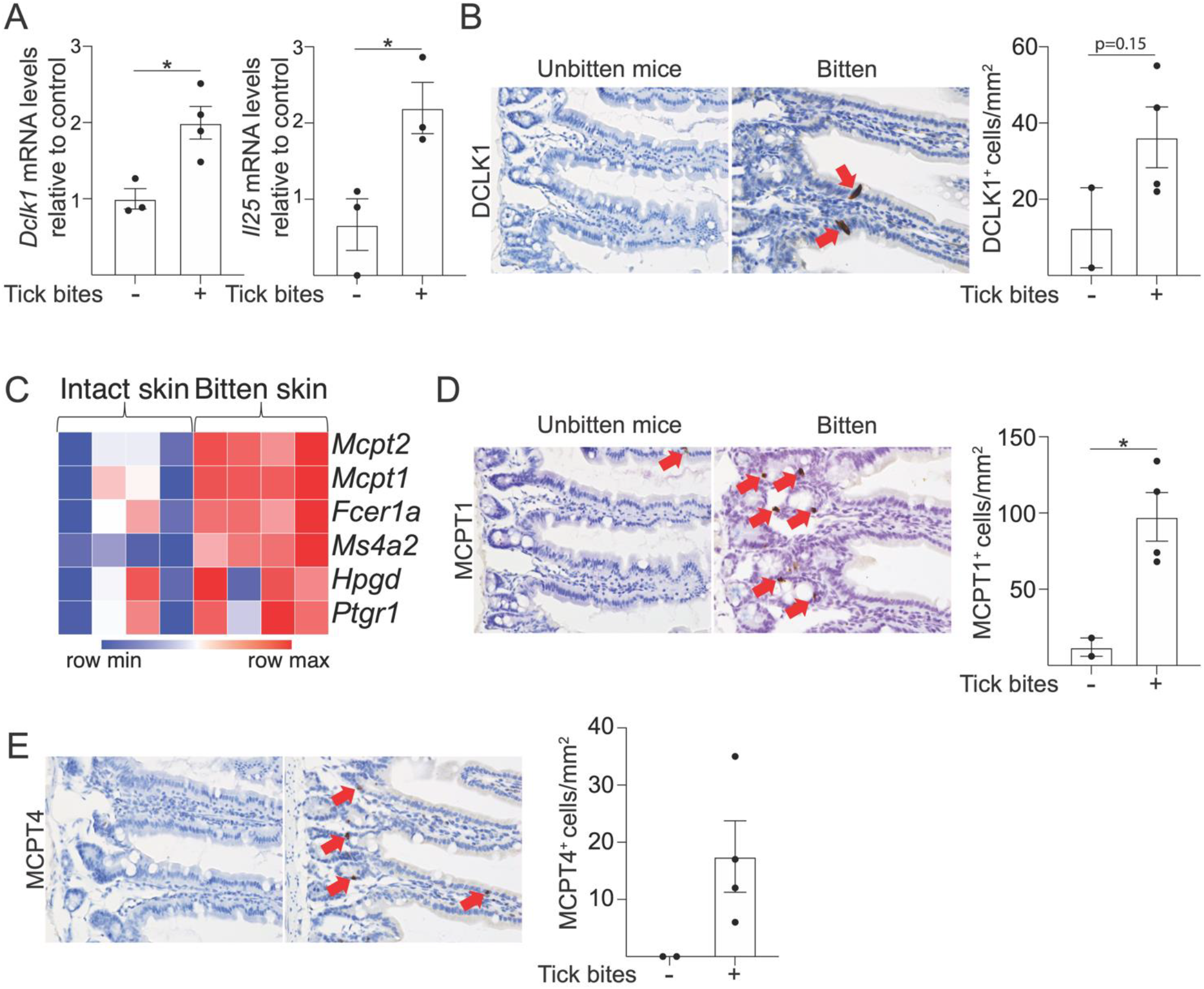
Tick bites from *Ixodes ricinus* promote an increase in the number of tuft cells and mast cells in the jejunum. **A-E**. Dclk1 (left) and Il25 (right) mRNA levels (A), representative DCLK1 immunohistochemistry staining and quantitation (B), heat map of mast cell gene signature (C), representative MCPT1 (D) and MCPT4 (E) immunohistochemistry staining and quantitation in the jejunum of unbitten or *I*. *ricinus* bitten on the skin of C3H/HEN mice at day 10. Red arrows indicate DCLK1+ (B), MCPT1+(D) and MCPT4+ (E) cells *p<0.05 by t test (one simple experiment with n=3-4 mice per experiment).

## DISCUSSION

Our results showed that *I. ricinus* bites in dorsal skin of C3H/HEN mice promote cutaneous inflammation resembling allergic skin inflammation, a gene expression profile similar to tape stripped skin and the expansion of jejunal tuft cells and mast cells. As most of the experiment were carried with field-collected ticks, we cannot exclude a potential role of microbiome or pathogens carried by the ticks in the responses observed in our study.

Our histological analyses are consistent with the histopathological observations detected in humans following tick bites, such as acanthosis, epidermal hyperplasia, eosinophil infiltration around blood vessels and into the hypodermis (3, 4). In addition, our results indicate that *I. ricinus* bites promote skin inflammation at day 10 which resembles histologically in some pathological features to atopic dermatitis lesional skin and seen in mouse models of this disease, such as epidermal hyperplasia, increased accumulation of eosinophils and mast cells and increased total IgE serum levels (28, 29).

Our results showed that *I. ricinus* bites and mechanical skin injury shared transcriptomic profiles and activate common pathways. Mechanical skin injury induced by tape stripping promotes the skewing of skin dendritic cells to favor Th2 over Th1 polarization (18, 26). Similarly, *I. ricinus* saliva injection enhances the capacity of dendritic cells to promote Th2 polarization but suppressed Th1 polarization (30). However, whether similar mechanisms are activated in the priming of the immune response in the skin by mechanical skin injury and tick bites to promote a type 2 dominated immune responses need further investigation.

Our results showed that *I. ricinus* bites promote the expansion of tuft cells, mucosal mast cells and connective tissue mast cells. We previously showed that mechanical skin injury induced by tape stripping promotes the expansion of intestinal mast cells in the jejunum by increasing the number of tuft cells, type 2 innate lymphoid cells (ILC2s) and by promoting their production of IL-4 locally in the jejunum (21). Mast cells expansion induced by tape stripping exacerbates passive and active food anaphylaxis. Whether a similar mechanism in the gut is activated by tick bites in the gut to promote mast cell expansion and if this expansion plays a role in red meat allergy remains to be investigated.

During the tick blood meal, tick-derived bioactive molecules are inoculated into the skin (14), including highly glycosylated proteins, such as vitellogenin and hemelipoglycoprotein (31). Remarkably, these glycosylated proteins cross-react with serum from alpha-gal syndrome patients (31). Whether glycosylated proteins from different tick species are mediating alpha-gal sensitization or reaching the gut to trigger active transport of red meat proteins with molecular mimicry needs further exploration.

## Author Contributions

**Conceptualization:** JML-C and NB, **Data curation, Formal analysis, and Visualization:** JML-C, **Investigation**: JML-C, MS, SEMS, DV-M, ME, PV, **Resources:** JC, **Supervision:** JML-C and NB, **Writing:** JML-C and NB

## Acknowledgments

J.M.L.C. was supported by Boston Children’s Hospital OFD/BTREC/CTREC. We thank Dana- Farber/Harvard Cancer Center in Boston, MA, for the service provided by the Rodent Histopathology Core. Dana-Farber/Harvard Cancer Center is supported in part by an NCI Cancer Center Support Grant # NIH 5 P30 CA06516. We also thank Dr. Olivier Rais (Institut de Zoologie, University of Neuchâtel, Switzerland) for providing *Ixodes ricinus* from a laboratory-reared colony, Dr. Elizabeth Jacobsen (Department of Immunology, Mayo Clinic Arizona) for providing MBP antibody and Dr. Hans Oettgen for reading the manuscript and useful discussions.

## Data Availability Statement

The data presented in the study are deposited in the Gene Expression Omnibus (GEO) repository, accession numbers GSE247060, GSE247064 and GSE247065.

